# Physics-informed machine learning for automatic model reduction in chemical reaction networks

**DOI:** 10.1101/2024.03.20.585845

**Authors:** Joseph Pateras, Colin Zhang, Shriya Majumdar, Ayush Pal, Preetam Ghosh

## Abstract

Physics-informed machine learning emerges as a transformative approach, bridging the gap between the high fidelity of mechanistic models and the adaptive, data-driven insights afforded by artificial intelligence and machine learning. In the realm of chemical reaction network modeling, this synergy is particularly valuable. It offers a solution to the pro-hibitive computational costs associated with detailed mechanistic models, while also capitalizing on the predictive power and flexibility of machine learning algorithms. This study exemplifies this innovative fusion by applying it to the critical biomedical challenge of A*β* fibril aggregation, shedding light on the mechanisms underlying Alzheimer’s disease. A corner-stone of this research is the introduction of an automatic reaction order model reduction framework, tailored to optimize the scale of reduced order kinetic models. This framework is not merely a technical enhancement; it represents a paradigm shift in how models are constructed and refined. By automatically determining the most appropriate level of detail for modeling reaction networks, our proposed approach significantly enhances the efficiency and accuracy of simulations. This is particularly crucial for systems like A*β* aggregation, where the precise characterization of nucleation and growth kinetics can provide insights into potential therapeutic targets. The potential generalizability of this automatic model reduction technique to other network models is a key highlight of this study. The methodology developed here has far-reaching implications, offering a scalable and adaptable tool for a wide range of applications beyond biomedical research. The ability to dynamically adjust model complexity in response to the specific demands of the system under study is a powerful asset. This flexibility ensures that the models remain both computationally feasible and scientifically relevant, capable of accommodating new data and evolving understandings of complex phenomena.

## 1 Introduction

Deriving models for complex processes is a hallmark of modern science. Cleverly devised modeling affords viability to the computational analysis of many complex physical, chemical, biological, geological, etc., systems. As system complexity grows, so too must the dimensionality of our models. Generally, the problem presented here belongs to a large class of multiphysics problems, which might be computationally difficult due to scale, underpinning dynamics, or observational burden. In the quest to unravel the complexities of Alzheimer’s disease (AD), particularly the pathological aggregation of Amyloid-*β* (A*β*) peptides, this study leverages advanced computational biology and machine learning methods to refine our understanding and modeling of A*β* fibril formation.

The aggregation of A*β* peptides, a central event in AD pathology characterized by the accumulation of insoluble fibrils detrimental to neuronal cells, poses a significant challenge due to the intricate biochemical processes involved, and the difficulty in observing them *in vivo*. Traditional modeling approaches, while valuable, often fall short in capturing the full dynamics of A*β* aggregation, necessitating novel strategies for model development and analysis. By employing physics-informed machine learning paradigms, this research aims to achieve automatic model order reduction. It focuses on accurately representing primary and secondary nucleation states: key processes in fibril aggregation involving the formation of nucleation centers and the generation of new nucleation sites on existing fibrils, respectively. Through the integration of accelerated and reliable parameter estimation techniques, our goal is to elucidate the nuanced dynamics of reduced order A*β* aggregation models, thereby contributing to the broader understanding of AD mechanisms and the exploration of potential therapeutic interventions.

In the rapidly evolving landscape of computational biology and machine learning, the integration of data-driven approaches with traditional scientific modeling has opened new vistas for understanding complex biological processes. One of the most pressing and enigmatic challenges in modern medicine is unraveling Alzheimer’s disease mechanisms, particularly the role of Amyloid-*β* (A*β*) fibril aggregation. This process, central to the pathology of AD, involves the accumulation and deposition of A*β* peptides into insoluble fibrils, which are toxic to neuronal cells. Despite extensive research, the precise dynamics and regulatory mechanisms of A*β* aggregation remain only partially understood, largely due to the complexity of the underlying biochemical processes and the limitations of conventional modeling techniques [1, 2]. While most common, regularly-forming forms of proteins are soluble, and are usually monomeric species or parcels of defined complexes, some functional amyloid states are insoluble, and as such pathological [3].

Ordinary differential equation modeling serves as a powerful tool in the study of amyloid-*β* fibril aggregation: a critical process implicated in the pathogenesis of Alzheimer’s disease. These models can elucidate key parameters such as nucleation rates, growth kinetics, and the influence of various environmental factors, ultimately contributing to a deeper understanding of the disease and the potential development of therapeutic interventions [4]. State-of-the-art analytical frameworks, such as Amylofit [5], utilize comprehensive rate equations to model the dynamics of filamentous assembly. These frameworks underscore three pivotal mechanisms influencing aggregation kinetics. First, primary nucleation is identified, a process solely influenced by the abundance of free monomers. An instance of this mechanism is the formation of nucleation centers in supersaturated solutions. Next, secondary nucleation is discussed, a phenomenon that is governed by both the availability of free monomers and the presence of pre-formed aggregate structures. An example of secondary nucleation is the formation of new nucleation sites on the surfaces of existing fibrils. Finally, the concept of fragmentation is introduced, a process that is independent of monomer concentration and relies solely on the existing aggregated mass. However, the relative importance of each process, and their modeling applicability are hyper-specific and depend upon the availability, veracity, and granularity of available data. As medical imaging, and the general observability of A*β* proteins is enabled further, the tasks of parsing new data and prescribing appropriate models must maintain pace. This work, presented with data on transient concentrations of A*β*, proposes a framework by which primary and secondary nucleation distinctions can be informatively drawn.

The challenge of accurately modeling complex biochemical systems—such as the aggregation of A*β* peptides into amyloid fibrils, a hallmark of Alzheimer’s disease, stems from their inherent multiphysics nature. These systems are characterized by intricate interactions across different physical, chemical, and biological domains. Traditional modeling techniques, while powerful, often struggle to capture the full spectrum of dynamics at play due to computational limitations and the difficulty of incorporating all relevant phenomena into a single model [6].

Thus, we propose a physics-informed neural network (PINN) model for amyloid-*β* parametric studies. This PINN architecture alleviates concerns in complex modeling through physics-informed biases, which allow learning paradigms to intuit relationships in incomplete, or noisy datasets, while increasing the speed and veracity of machine learning models [7, 8]. Physics-informed machine learning emerges as a cutting-edge solution to the limitations faced by conventional modeling approaches [9, 11], offering a novel pathway to model complex systems like A*β* aggregation with unprecedented accuracy and efficiency where hidden physics in complex multiphysics problems can be intuited by learning algorithms [12, 13].

In the context of A*β* aggregation, PINNs represent a significant leap forward. One of the most compelling advantages of PINNs is their ability to overcome the computational and data-related challenges that hamper traditional modeling efforts. Ultimately, the integration of physics-informed machine learning into the study of A*β* aggregation aims to catalyze a deeper understanding of Alzheimer’s Disease pathogenesis, promising to accelerate the development of effective treatments [14].

By incorporating domain knowledge of chemical kinetics and reaction dynamics directly into the learning algorithms, we aim to develop a model that not only predicts the behavior of A*β* aggregation with high accuracy but also ad-heres to the well-studied biophysical principles governing these processes. This approach enables the exploration of the aggregation phenomenon beyond the constraints of traditional computational models, offering insights into the kinetics of nucleation, growth, and the formation of toxic fibrils under varying conditions.

## 2 Methods

### 2.1 Reduced order chemical kinetic model

The uncontrolled aggregation of toxic amyloid fibrils is a condition underpinning many serious health conditions. An example of such a protein, which reacts with environmental catalysts in our brains, is called Amyloid-*β* (A*β*). Modeling the formation of potentially toxic amyloid plaques is important [15] as direct observation of this process is prohibitively invasive. In the year 1906, it was only during an autopsy when Dr. Alois Alzheimer noted the presence of these plaques in the brain of a patient exhibiting dementia like symptoms now synonymous with Alzheimer’s Disease.

The model presented in figure 6 (in Appendix A) is a reduced order representation of A*β* aggregation. Reduced order representations are studied thoroughly to draw conclusions about competing aggregation processes [16, 17]. The entire amyloid aggregation system is complex and a full-scale analysis is difficult to imagine. The aggregation of A*β* into amyloid fibrils involves a series of bio-chemical reactions, including nucleation, elongation, and fibril branching. The full-scale simulation of every individual reaction and molecular interaction is computationally infeasible due to the sheer number of molecules and the stochastic nature of their interactions. Therefore, reduced order models are essential for understanding the key processes that drive A*β* aggregation. These models abstract the aggregation process into a series of rate equations that describe the change in concentration of different species over time, focusing on critical transitions such as nucleation and fibril elongation.

Reduced order models are derived based on the assumption that certain steps in the aggregation process are rate-limiting, meaning they significantly affect the overall rate of aggregation. For example, the formation of a stable nucleus (the nucleation process) is often considered a critical step in the aggregation pathway. By identifying and modeling these key steps, researchers can gain insights into the mechanisms of aggregation and identify potential thera-peutic targets to prevent or disrupt the formation of toxic amyloid fibrils. The mass-action framework for constructing the reduced order representation of the aggregation complex is given in Appendix A, where the On- and Off-pathway toward aggregation are considered.

### 2.2 PINN for parameter estimation

Machine learning approaches are popular for their ability to transcend many of the costs associated with large-scale models. Machine learning in multiphysics systems can explore high dimensional feature spaces. Deep architectures offer creative ways to extract features from multi-fidelity data. Traditional machine learning algorithms, such as neural networks, are data-driven and rely solely on input-output relationships observed in the data. While these models can perform well when large and representative datasets are available, they might struggle when data is limited, noisy, or subject to physical constraints.

Some problems lack the massive empirical data to train a model or are prohibited by the computational expense of reliable simulation data. Physics-informed machine learning harnesses the computational advantages of learning machines, while accelerating training and improving model generalization by integrating the model with systemic information. Karniadakis et al. [9] describe three principles of physics-informed machine learning: observational biases, inductive biases, and learning biases. The proposed PINN model for chemical reaction networks draws from learning biases informed by governing equations and inductive bias informed by underlying knowledge of the aggregation kinetics. As categorized by the drivers described in Pateras et al. taxonomy of PIML drivers and biases [10], governing kinetics and underlying subject area expertise inform model architecture, learning processes and binning choices: providing physics-model and physics-data driven learning and inductive biases.

By incorporating the known governing equations as constraints during the training process, physics-informed machine learning can effectively estimate the final outcome of the aggregation process, and the time-dependent effect of seedings. Physics-informed machine learning in partial differential equations has shown promising results in various domains and has the potential to accelerate scientific discovery and engineering design by efficiently identifying parameters and providing accurate solutions to complex physical systems, with extensions in both labelled and unsupervised learning. The integration of known governing equations as constraints ensures that the learned parameters satisfy the underlying physical laws, making the predictions more reliable and interpretable. For instance, in fluid dynamics, heat transfer, materials science, and many other fields, this technique has facilitated efficient and accurate estimation of parameters that govern the system’s behavior. PINNs represent a fusion of data-driven learning and physics-based constraints, enabling the estimation of model parameters that are not only consistent with observed data but also adhere to the fundamental laws of physics. This integration is crucial for modeling systems like A*β* aggregation, where experimental data might be sparse or incomplete, and the biological processes are governed by complex biochemical reactions. PINNs offer a computationally efficient alternative to traditional simulation methods, particularly when dealing with complex multiphysics systems like protein aggregation. By training the neural network to satisfy both the data and the physics-based constraints, PINNs can bypass the need for extensive simulations, reducing the computational cost and time required for parameter estimation and model validation.

Toward further expanding these fibrillar studies, a physics-informed neural network, infused with model-driven learning bias, is employed to predict time series of amyloid-*β* concentrations. As with the observational bias model, a neural network performs the task of regression. However, rather than just predicting final aggregation outcomes, the learning bias model is able to predict the time-dependent effect of aggregate seeding. In the context of A*β* aggregation, where direct measurements of intermediate species concentrations over time can be challenging to obtain, PINNs offer a powerful tool for extrapolating from limited data. By combining sparse observational data with the robust framework of the aggregation model’s differential equations, PINNs can provide detailed insights into the kinetics of the aggregation process, including the effects of nucleation, growth, and fibril formation.

Figure 1 gives a schematic of the learning bias processed. Here the physics-informed loss is jointly optimized with neural network loss. Equations 1 - 2 describe the different computations performed. Physics-informed loss, ℒ_*PI*_, forces the model to adhere to governing physical principles, while standard neural network loss, ℒ_*MSE*_, directly learns correlations in training data. By embedding the governing equations of A*β* aggregation directly into the loss function of the neural network, PINNs can predict the dynamics of amyloid formation under various conditions. This method ensures that the predictions respect conservation laws, reaction kinetics, and other biophysical constraints, enhancing the model’s reliability and interpretability. The physics-informed loss component ℒ_*PI*_ penalizes deviations from the physical model, guiding the learning process to conform to the biophysical reality of amyloid fibril formation. Here, *F* is the numerically derived solution to the governing equations, while *Ŷ*_*t*_ and *Y*_*t*_ denote neural network predicted labels and training observations respectively. Equation 2 forces predictions to match training data, while 1 forces predictions to match the underlying dynamics. Thus, aggregate loss ℒ is defined as ℒ = ℒ_*MSE*_ +ℒ_*PI*_ .

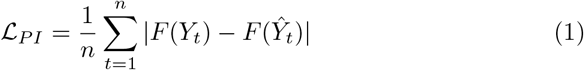

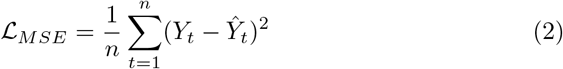

**Figure 1.**
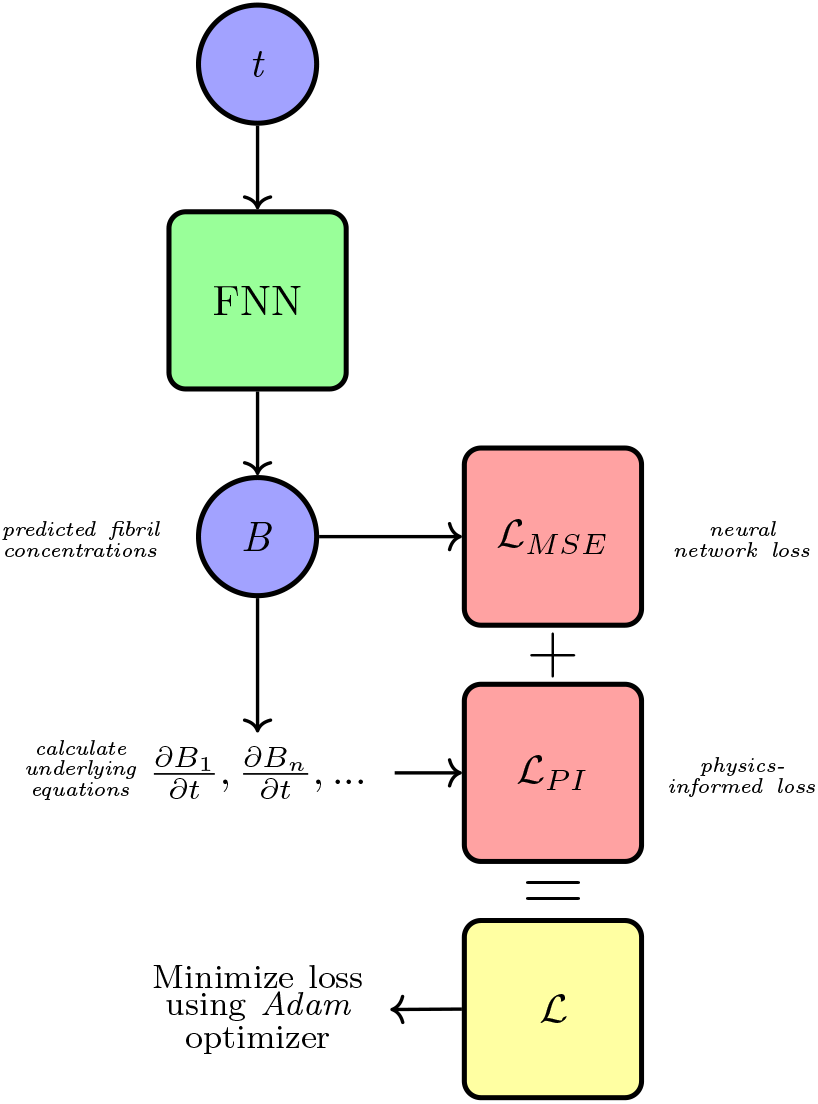
A diagram of physics-informed learning bias in the informed neural network routine. Observed aggregation is used to generate loss from data and from governing equations.

Incorporating PINNs into the study of A*β* aggregation offers a novel approach to understanding and predicting the dynamics of amyloid fibril formation. This methodology not only provides a pathway to estimate unknown parameters with high accuracy but also opens new avenues for exploring the mechanistic underpinnings of neurodegenerative diseases. Through the synergistic combination of machine learning and biophysical modeling, researchers can uncover new insights into the complex process of amyloid aggregation, paving the way for the development of more effective diagnostic and therapeutic strategies.

### 2.3 Automatic reaction order reduction

We formalize the proposed automatic model reduction framework by answering the following question: **Given data on amyloid concentration of several molecular weight distinctions, what is the optimal choice of modeling scale for primary and secondary nucleation species toward reliably balancing model reduction for complexity and model granularity?**

Given the influence of nucleation in rate limiting the aggregation process, it is recognized the model must differentiate between pre-nucleation and post-nucleation events to capture the essential physics of A*β* aggregation. The process of nucleation is a cornerstone in the aggregation of A*β*, serving as a rate-limiting step that significantly influences the overall dynamics of aggregation. The distinction between primary-nucleation and secondary-nucleation events is not merely a technical detail but a fundamental aspect that affects how models predict the formation and growth of amyloid fibrils. Pre-nucleation interactions lead to the formation of oligomers, which then transition through a nucleation event to form the seeds for fibril elongation. Differentiating between these stages is crucial for understanding how initial conditions and environmental factors influence the aggregation pathway and the formation of toxic versus non-toxic species. The accurate modeling of these stages is critical for understanding the initiation and progression of amyloid diseases, making the choice of modeling scale a question of paramount importance. Thus by harnessing domain specific knowledge about nucleation and utilizing learning machines which facilitate the fusion of physical priors and optimization algorithms, we can answer the question presented above with domain-specific intuition and with learning machine optimality.

As explored in Appendix A, the 3-species on-pathway model has been studied for certain datasets. Reduced order models for ordered aggregation of A*β* are constructed based on assumptions that identify the rate-limiting steps both before and after nucleation, which are critical in the formation and aggregation process of amyloid fibrils. These models simplify the complex mechanisms of amyloid beta aggregation by focusing on key transitional states, such as the formation of oligomers and their assembly into fibrils, underpinning the aggregation pathway. The efficacy of these models in extrapolating off-pathway aggregation and amyloidosis lies in their ability to predict the behavior of A*β* under various conditions. By focusing on rate-limiting steps, these models help in understanding how deviations from the primary pathway lead to disease-relevant off-pathway aggregates, offering a valuable tool for the development of strategies to combat amyloid-related diseases.

When presented with a novel dataset, we propose a method for optimal model scale reduction. For example: consider a dataset with measurements of monomer, pre-nucleation oligomers, and post-nucleation fibrils. Where each measurement is marginally close enough to each neighbor to consider model reduction in either direction. Figure 2 serves as a visual guide, illustrating the possible pathways for model reduction and the decision-making process involved in selecting an optimal scale. This diagram underscores the iterative nature of the process, where each step represents a critical decision point in achieving a balance between model simplicity and the need to capture essential dynamics. The development of automatic network models for A*β* aggregation is urgent for several reasons:

**Figure 2.**
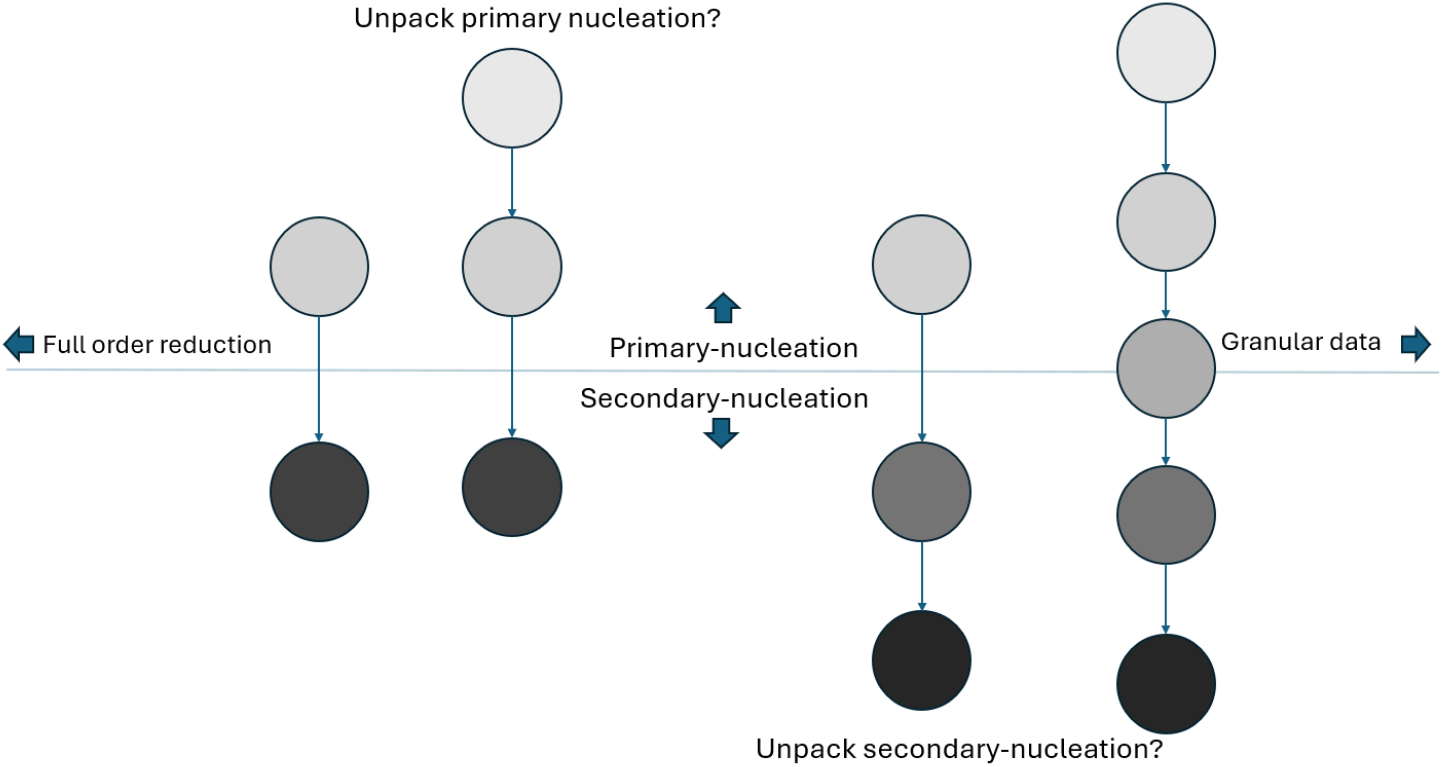
A diagram of the on-pathway model reduction choice confronting the problem of optimal reduced order scaling.

1. Complexity vs. Reliability Trade-off: As the complexity of the biological system increases, so does the challenge of capturing all relevant dynamics without oversimplifying the system. Automatic network models offer a pathway to navigate this trade-off, enabling the selection of a modeling scale that retains essential features while remaining computationally feasible.
2. Advancements in Treatment Strategies: Understanding the detailed mechanisms of A*β* aggregation is essential for developing effective treatment strategies for Alzheimer’s disease. Automatic network models can accelerate this process by providing insights into how changes at the molecular level influence the formation of toxic fibrils.
3. Adaptability to New Data: The field of amyloid research is rapidly evolving, with new data constantly emerging. Automatic models that can adapt to new datasets and refine their parameters accordingly are crucial for keeping pace with advancements in the field.

The proposed workflow for automatic reaction order reduction represents a methodological innovation that addresses these challenges. By iteratively fitting the Reduced Order Model Physics-Informed Neural Network (ROMPINN) across different scales of model reduction and assessing the fit, researchers can identify the most appropriate level of granularity for their model. This approach not only enhances the model’s reliability but also its relevance to current and future datasets. The informed reaction unpacking for automatic network model selection represents a potential significant step forward in the field, offering a systematic approach to tailoring models to the specific dynamics of amyloid aggregation. This methodological innovation not only advances our understanding of Alzheimer’s disease but also opens new avenues for the development of ther-apeutic interventions, illustrating the profound impact of modeling techniques on the fight against neurodegenerative diseases.

The workflow for our proposed model reduction framework is as follows:

1. Fit ROMPINN for maximally reduced order model. Return residuals of fit.
2. Propose crucial primary rate-limiting steps by unpacking primary-nucleation reactions, rearrange governing reactions, and fit. Explore parameter identifiability. Backtrack without improvement.
3. Propose crucial secondary rate-limiting steps by unpacking secondary-nucleation reactions, rearrange governing reactions, and fit. Explore parameter identifiability. Backtrack without improvement.

Eight potential model architectures are explored in this study, each coalescing or unpacking primary and/or secondary nucleation reactions in unique ways. Figure 3 depicts the eight architectures, where Model 8 is displayed with its corresponding forward and backward reaction parameters, *α* and *β* respectively.

**Figure 3.**
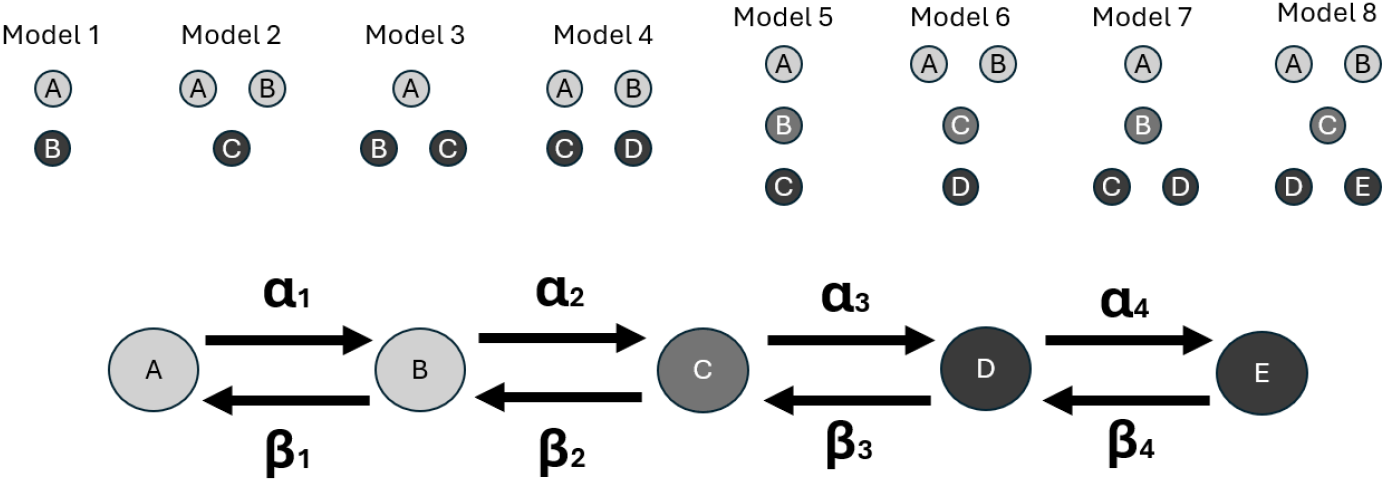
A diagram of the on-pathway model reduction choice confronting the problem of optimal reduced order scaling. Fully granular Model 8 is displayed.

This model reduction approach explores the model architecture space in a limited scope – adding or removing granularity for primary or secondary nucleation species. Future work must undoubtedly include exploration of other important rate limiting factors toward pathogenesis. The aggregation process encompasses several biophysical processes that contribute to the formation and accumulation of A*β* fibrils, which are central to the disease. Several other models are explored employing Amylofit, and further reinforce the need to include other subject area expertise in the generic model reduction architecture. We briefly summarize these models in the following.

First, nucleation and elongation model posits that A*β* aggregation begins with a nucleation phase, where monomers come together to form a stable nucleus, followed by an elongation phase, where additional monomers add to the growing fibril. The nucleation phase is considered a rate limiting step because it requires overcoming an energetic barrier. Once a nucleus is formed, the elongation phase can proceed more rapidly. This model is important for understanding the initial stages of amyloid fibril formation and highlights the significance of targeting early nucleation events in therapeutic interventions.

In the secondary nucleation dominated model, the emphasis is on the formation of new nuclei on the surfaces of existing fibrils. This process significantly accelerates the aggregation of A*β*, leading to a faster accumulation of amy-loid fibrils. Secondary nucleation is a key mechanism by which small changes in the concentration of A*β* can lead to dramatic increases in fibril formation, underlining its importance in the exponential phase of amyloid accumulation.

The fragmentation dominated model highlights the role of fibril fragmentation in generating new seeds for further aggregation. Fragmentation increases the number of active ends, facilitating the addition of monomers and accelerating fibril formation. This process is crucial for understanding how fibril breakage contributes to the spread and severity of amyloidosis.

The fragmentation and secondary nucleation model combines aspects of both fragmentation and secondary nucleation, recognizing that both processes can significantly contribute to A*β* aggregation. It suggests a synergistic effect where fragmentation creates new ends that enhance elongation, while secondary nucleation increases the number of nucleation sites. This dual mechanism provides a more comprehensive understanding of how A*β* fibrils proliferate and accumulate. Finally, the multistep secondary nucleation dominated model is considered.

This this model proposes that secondary nucleation involves multiple steps, including the attachment of A*β* monomers to fibril surfaces, structural changes to form a nucleus, and the eventual detachment of new nuclei to seed further fibril formation. It highlights the complexity of the nucleation process and suggests multiple potential targets for therapeutic intervention. This model underscores the intricate balance and interplay between different biophysical processes in amyloidosis and the importance of targeting specific steps in the aggregation pathway to combat diseases like Alzheimer’s.

## 3 Results

First, we examine the utility of ROMPINN to fit to observed data with reliable residuals. All eight reduced order binning structures are capable of fitting data with mean absolute error *<* 0.08. Figure 4 displays the physics-informed and data driven cumulative training losses, while table 1 displays the comparison between residuals fit with the ROMPINN exploratory architecture and state-of-the-art amyloid system models [5]. While mean residual values for ROMPINN are relatively consistent, the utility of models for accurate forecasting is dependent more on the binning structure and ability to fit data. The balance of fit residual, forecasting ability, and parameter identifiability is what affords a proposed model architecture *optimality* or, rather, make it the current best choice for order reduction in relation to the available observations. Identifiability for each parameter are calculated employing COPASI parameter scans [18] and the process is detailed in Appendix B. The profile likelihood method is a statistical technique used to estimate the confidence intervals for parameters within complex models, especially when dealing with multiple parameters or constrained estimation problems. It operates by fixing the parameter of interest at various values while optimizing the likelihood function with respect to all other estimable parameters, thereby constructing a profile of the likelihood as a function of the parameter of interest. This profiling process allows for a more accurate estimation of the confidence interval for the parameter by taking into account the uncertainty and interdependence of all model parameters.

**Table 1:** Comparison between ROMPINN and state-of-the-art amyloid fitting tools.

| Model | Mean Residual |
| --- | --- |
| Levenberg-Marquardt: |  |
| Model 8 | 0.10912 |
| ROMPINN: |  |
| Model 1 | 0.054556 |
| Model 2 | 0.016991 |
| Model 3 | 0.021339 |
| Model 4 | 0.055874 |
| Model 5 | 0.020794 |
| Model 6 | 0.075503 |
| Model 7 | 0.080127 |
| Model 8 | 0.060492 |
| Amylofit: |  |
| nucleation, elongation | 0.0703 |
| secondary nucleation dominated | 0.001019 |
| fragmentation dominated | 0.000489 |
| fragmentation and secondary nucleation | 0.000913 |
| multistep secondary nucleation dominated | 0.057 |

**Figure 4.**
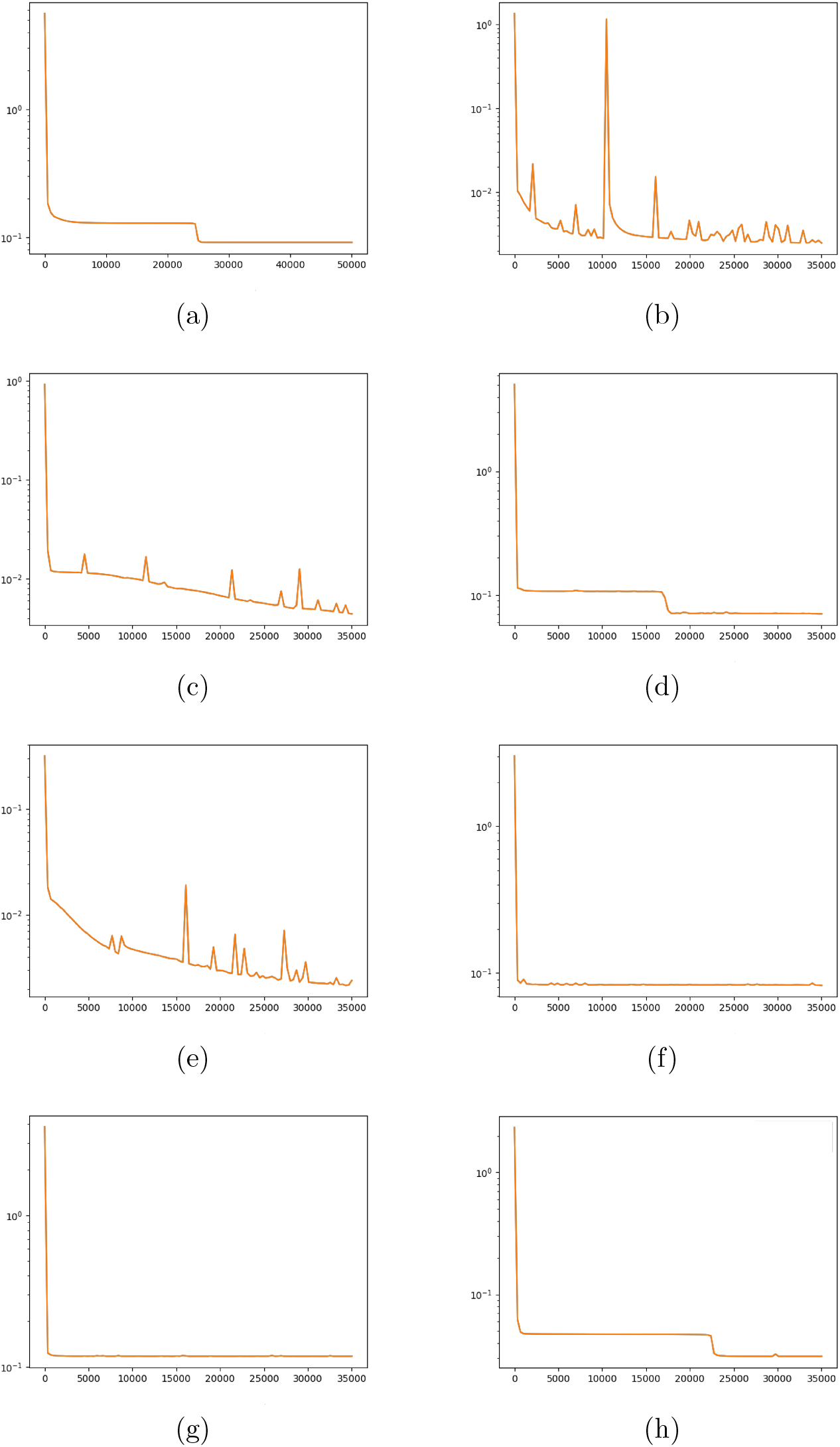
The cumulative physics-informed and observational training loss for (a) Model 1, (b) Model 2, (c) Model 3, (d) Model 4, (e) Model 5, (f) Model 6, (g) Model 7, and (h) Model 8.

ROMPINN is generally capable at producing reliable fits to data - as displayed in Table 1 - while showcasing the dual purpose of the PINN module for reaction parameter estimation in Table 2. ROMPINN generally produces tighter fit to observed data than traditional least squares methods. State-of-the-art methods particularly fine-tuned for A*β* aggregation modeling outperform in terms of fit to transient concentrations [5]. However the generalizability off the ordered aggregation model, coupled with reliably identifiable parameters, allow for further lines of inquiry for seeding studies, reaction network thermodynamic spontaneity, and stability of off-pathways toward particularly toxic pathogenic states [16].

**Table 2:**
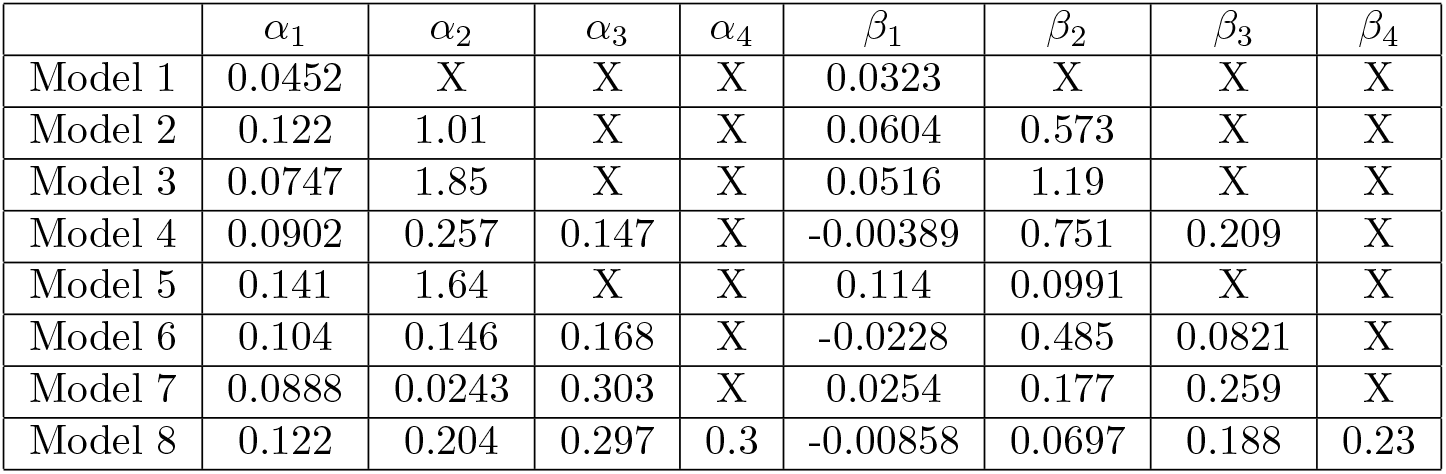
Reaction rate parameters estimated by ROMPINN routine.

| | $\alpha_1$ | $\alpha_2$ | $\alpha_3$ | $\alpha_4$ | $\beta_1$ | $\beta_2$ | $\beta_3$ | $\beta_4$ |
| --- | --- | --- | --- | --- | --- | --- | --- | --- |
| Model 1 | 0.0452 | X | X | X | 0.0323 | X | X | X |
| Model 2 | 0.122 | 1.01 | X | X | 0.0604 | 0.573 | X | X |
| Model 3 | 0.0747 | 1.85 | X | X | 0.0516 | 1.19 | X | X |
| Model 4 | 0.0902 | 0.257 | 0.147 | X | -0.00389 | 0.751 | 0.209 | X |
| Model 5 | 0.141 | 1.64 | X | X | 0.114 | 0.0991 | X | X |
| Model 6 | 0.104 | 0.146 | 0.168 | X | -0.0228 | 0.485 | 0.0821 | X |
| Model 7 | 0.0888 | 0.0243 | 0.303 | X | 0.0254 | 0.177 | 0.259 | X |
| Model 8 | 0.122 | 0.204 | 0.297 | 0.3 | -0.00858 | 0.0697 | 0.188 | 0.23 |

Table 2 details the estimated parameters from each proposed model architecture. Note when binning is done fully, only one reversible reaction remains: where *α* and *β* represents forward and backward rates respectively. However, the fully unpacked model contains eight parameters to be estimated.

Generally, employing parameter scans of the ordered aggregation network with Copasi parameter scans, [18] the parameters of Model 7 are mostly robust toward large permutations of the conditions. Likelihood profiles in Figure 5 reflect generally acceptable symmetry about the maximum likelihood estimate in reaction rates under scan: suggesting those rates are significantly determining steps in the reaction network. The task of comparing model architecture, PINN fit to data, and parameter identifiability suggest Model 7 the most apt - agreeing with the well understood importance of secondary nucleation reactions, while nodding toward the possibility of other important post-nucleation steps like fragmentation. Profile likelihoods with tight 95% confidence intervals about the maximum likelihood estimates suggest individually identifiable parameters. The relatively better identifiabilty in backwards, secondary nucleation reaction steps further suggests an important underlying rate-limiting process: further providing concordance with the understanding about the importance of secondary nucleation and fragmentation interactions.

**Figure 5.**
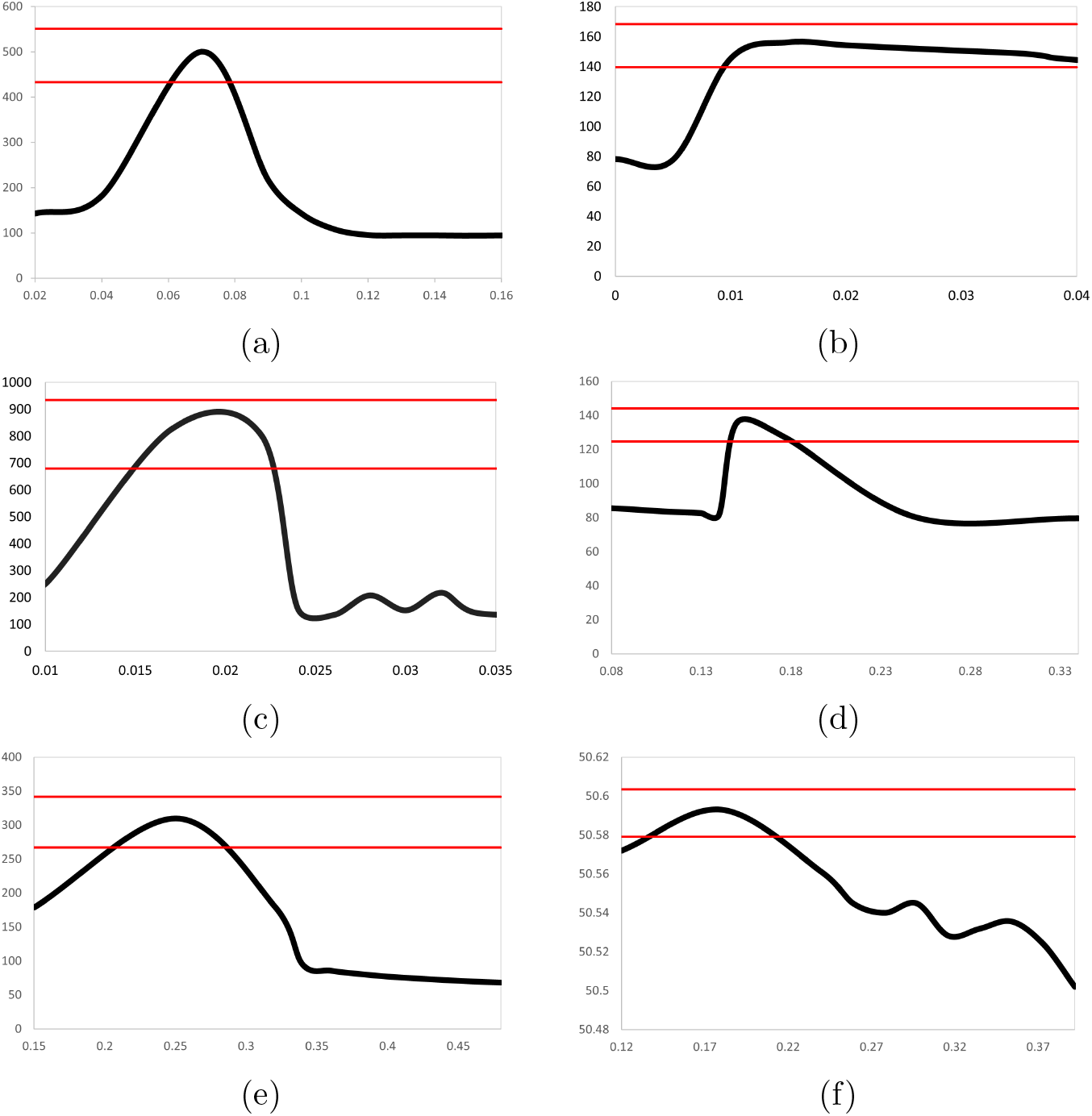
Likelihood profile for parameters estimated in Model 7. Profile like-lihood as a function of estimable parameters *αi* (a),(c),(e) and *βi* (b),(d),(f) where i = 1, 2 and 3. 95% confidence intervals displayed about the maximum.

**Figure 6.**
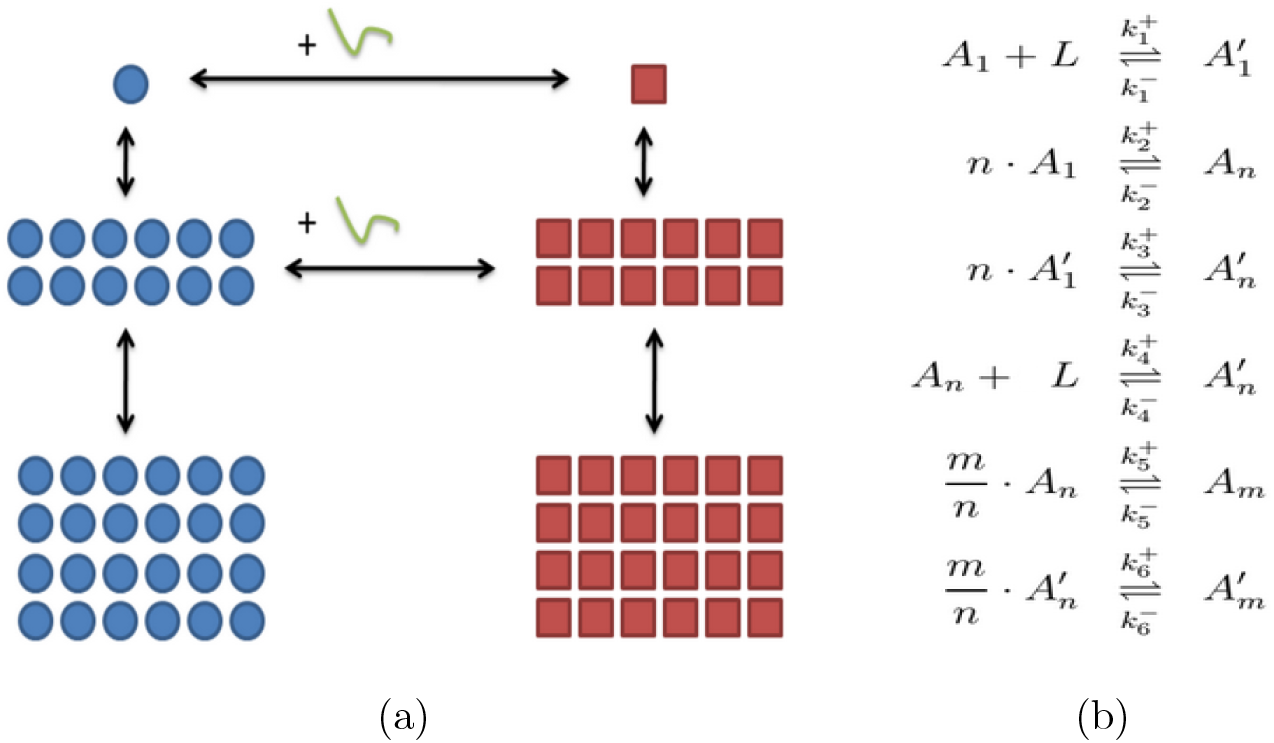
(a) A reduced order aggregation model for A*β* amyloid and (b) the chemical reactions corresponding to the link between each aggregation step or pathway mutation.

## 4 Discussion

The major contribution of this work is the development and application of the ROMPINN architecture, an approach in computational biology that leverages the strengths of machine learning, physics-informed modeling, and domain-specific knowledge to tackle the challenges posed by the complex biochemical processes underlying A*β* aggregation. This approach allows a more nuanced understanding of the dynamics of amyloid fibril formation, particularly differentiating between primary and secondary nucleation processes. By integrating accelerated and reliable parameter estimation techniques, this research advances our comprehension of AD mechanisms, potentially paving the way for novel therapeutic interventions and provides a framework for approaching novel data. The integration of parameter identifiability, PINNs and subject area expertise signifies a substantial advancement, in general, for modeling complex biological systems. This methodology not only enhances the accuracy and efficiency of the models but also ensures that they are grounded in biophysical reality.

Despite its notable contributions, this study is not without its limitations. One of the main restrictions is the massive utility of Amylofit, a state-of-the-art analytical framework used to model A*β* aggregation kinetics. While Amylofit provides a comprehensive understanding of filamentous assembly dynamics, it may not capture all the nuances of A*β* aggregation, particularly in the context of diverse biological environments and varying conditions. Furthermore, the ROMPINN architecture, though powerful, relies on the availability and quality of data for training and validation. The inherent complexity of A*β* aggregation and the stochastic nature of biochemical processes pose significant challenges to modeling efforts, requiring further refinement of the ROMPINN approach to fully capture the dynamics of the disease.

Future research should focus on several key areas to build upon the foundations laid by this study. Firstly, enhancing the data collection and processing methods to include a wider range of biological conditions and variables can improve the model’s robustness and applicability to other real-world scenarios. Expanding the ROMPINN architecture to incorporate other aspects of AD pathology, beyond A*β* aggregation, could offer more comprehensive insights into the disease mechanisms. Further exploration into the integration of multi-scale modeling techniques could also prove beneficial. This would allow for the bridging of molecular-level interactions with larger-scale pathological outcomes, offering a more holistic view of AD progression. Additionally, the exploration of novel machine learning algorithms and training methodologies could further refine parameter estimation techniques, enhancing the model’s predictive capabilities.

Another critical area for exploration involves leveraging the ROMPINN architecture’s capacity for studying off-pathway aggregation models - as fully discussed in the reduced order model of Appendix A. The ease of applicability of the framework to model ordered aggregation processes uniquely positions it to investigate hidden reaction kinetics, a dimension where traditional models like Amylofit may not fully venture. This capability is particularly valuable for uncovering the subtle, yet crucial, biochemical pathways that contribute to A*β* aggregation but remain obscured in conventional studies. Understanding these off-pathway reactions is essential for a comprehensive picture of A*β* aggregation dynamics, as they can significantly influence the rate, extent, and morphology of fibril formation. These pathways often involve intermediate species or alternative folding patterns that could potentially serve as therapeutic targets. By enabling the detailed study of such kinetics, the ROMPINN architecture can uncover novel insights into the aggregation process, further distinguishing this approach from existing models.

The exploration of hidden reaction kinetics through the application of ROMPINN to off-pathway aggregation models not only enhances our understanding of Alzheimer’s disease at a molecular level but also opens the door to identifying novel biomarkers and therapeutic targets. This future direction promises to expand the scope of computational biology in AD research, advancing beyond the limitations of current analytical frameworks to explore the full complexity of the disease. Moreover, the ROMPINN framework can be generalized to other chemical reaction networks where intricate rate limiting dynamics of the physical system can be unraveled leading to the generation of testable hypotheses.

## A Reduced order chemical kinetic model

The model in figure 6 describes individual monomers of A*β* aggregating to form a nucleation size oligomer of size *n*, and an ordered step of aggregation to form a full fibril containing m-many individual A*β* proteins [19]. *A*_1_ is the monomer species of A*β. n*-many *A*_1_ proteins come together to form an experimentally observed intermediate nucleation size, *A*_*n*_. Each *A* also has a corresponding ‘prime’ species, *A*’, signifying an aggregation reaction has occurred in the presence of an environmental catalyst *L*, such as a fatty acid or a surfactant. The inclusion of ’prime’ species (*A*′) in the model represents the impact of environmental catalysts, such as metal ions or fatty acids, on the aggregation process. These catalysts can alter the aggregation pathway [20], leading to the formation of structurally different oligomers and fibrils that may have distinct toxicological properties. By modeling the interactions between A*β* and these catalysts, the model can explore how changes in the biochemical environment influence the balance between healthy and toxic pathways. This differentiation is crucial for understanding the conditions under which A*β* aggregation becomes pathological, offering insights into potential strategies for disease intervention. Both healthy and toxic pathways culminate in a post-nucleation size oligomer of size *m* (*A*_*m*_ and 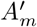 respectively). The reactions in figure 6(b) describe the interactions between the differently-sized aggregates of A*β*. Each double ended arrow in figure 6 describes a reversible aggregation reaction, or a switching between toxic and healthy pathways. It is observed that fully realized fibrils no longer mutate between healthy or toxic.

Each arrow in our network abstraction corresponds to a coalescing or dis-solving aggregation reaction. Employing the law of mass action we can convert the list of chemical reactions into a system of differential equations describing the concentration of each Amyloid-*β* species.

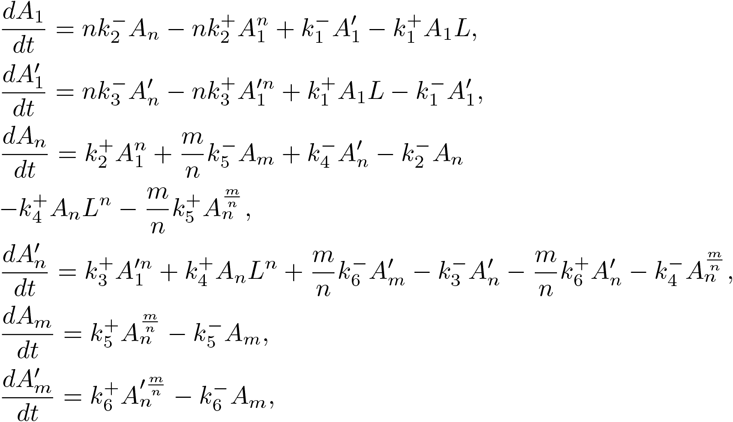

The reaction constants are defined:

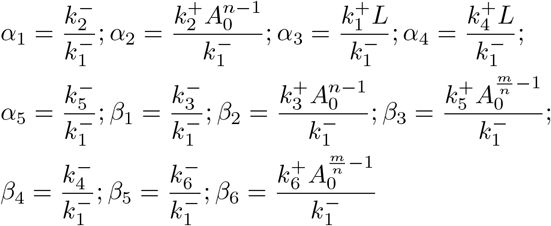

This system is then non-dimensionalized. We use *A*_0_ be the characteristic concentration of monomers and 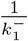 as the characteristic time, we define the dimensionless species as follows:

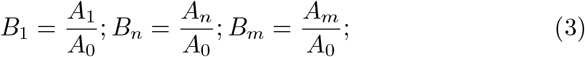

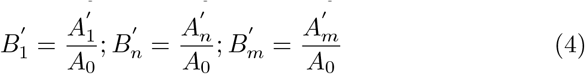

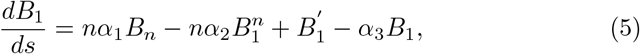

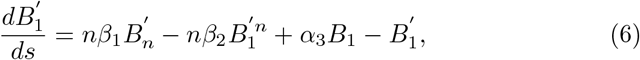

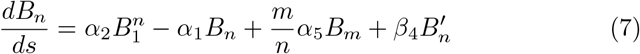

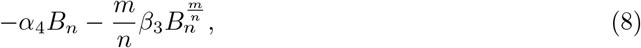

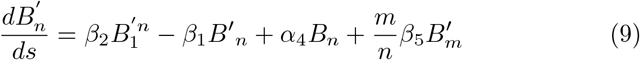

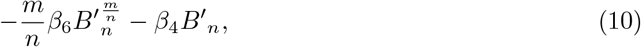

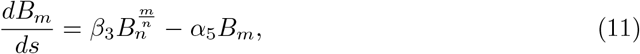

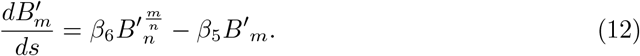

Here, *B*_1_, *B*_*n*_, *B*_*m*_ represent healthy oligomers of sizes 1, *n, m* and 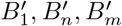 are toxic oligomers. The various nondimensional rate constants are given by *α*_*i*_ or *β*_*i*_. Time dependent concentration of A*β* plaques, derived by estimating solutions to equations 5 - 12, allow conclusions about how environmental conditions affect the aggregation process. The computation of the governing equations is often performed repeatedly for various initial conditions and for differing values of rate constants representing environmental factors.

Later, upon utilizing the reduced order model of amyloid aggregation for automatic network scaling, each proposed reduced order architecture obeys the same derivation of governing equations.

## B Parameter Identifiability

The likelihood function for a set of parameters *θ* (of interest) and *ψ* (nuisance parameters) based on observed data *X* is given by:

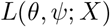

The profile likelihood for *θ* is defined by maximizing the likelihood function over the nuisance parameters *ψ* for each fixed value of *θ*:

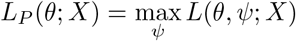

To estimate a confidence interval for *θ*, we use the profile likelihood ratio:

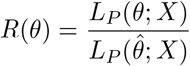

where 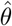 is the maximum likelihood estimate of *θ*, obtained by maximizing *L*_*P*_ (*θ*; *X*) over *θ*.

A 95% confidence interval for *θ* can then be approximated by finding the values of *θ* for which *R*(*θ*) exceeds a critical value, typically based on the chi-square distribution with degrees of freedom equal to the number of parameters of interest:

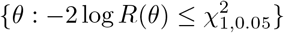

- *L*(*θ, ψ*; *X*) is the likelihood function for the model parameters given the data.
- *L*_*P*_ (*θ*; *X*) = max_*ψ*_ *L*(*θ, ψ*; *X*) is the profile likelihood for *θ*, obtained by maximizing over *ψ*.
- 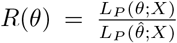 is the profile likelihood ratio, used to test hypotheses about *θ*.
- 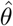 represents the maximum likelihood estimate of *θ*.

The confidence interval is determined by the set of *θ* values for which the profile likelihood ratio statistic −2 log *R*(*θ*) does not exceed the critical value from the chi-square distribution, 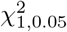, indicating a 95% confidence level.

This method adjusts the likelihood function to account for the uncertainty in the nuisance parameters, providing a more accurate and robust estimation of the confidence interval for the parameter of interest. In our COPASI parameter scan, the likelihood profile *L*_*P*_ (*θ*_*i*_) for each fitted parameter 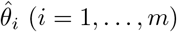 is defined as:

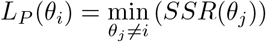

i.e., re-optimizing the objective function value *SSR*(*θ*_*j*_) with respect to all parameters *θ*_*j*_≠*i* for defined values of *θ*_*i*_ in a neighborhood of the original estimated parameter value 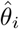. The parameter 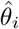 is called identifiable, if the reoptimized *SSR*(*θ*_*j*_) exceeds a certain confidence limit within the tested interval.

## References

[1] Hardy, J., & Selkoe, D. J. (2002). The amyloid hypothesis of Alzheimer’s disease: progress and problems on the road to therapeutics. Science, 297(5580), 353–356.

[2] Sadigh-Eteghad, S., Sabermarouf, B., Majdi, A., Talebi, M., Farhoudi, M., & Mahmoudi, J. (2015). Amyloid-beta: a crucial factor in Alzheimer’s disease. Medical principles and practice, 24(1), 1–10.

[3] Knowles, T. P., Vendruscolo, M., & Dobson, C. M. (2014). The amyloid state and its association with protein misfolding diseases. Nature reviews Molecular cell biology, 15(6), 384–396.

[4] Iwatsubo, T., Odaka, A., Suzuki, N., Mizusawa, H., Nukina, N., & Ihara, Y. (1994). Visualization of Aβ42(43) and Aβ40 in senile plaques with end-specific Aβ monoclonals: Evidence that an initially deposited species is Aβ42(43). Neuron, 13(1), 45–53.

[5] Meisl, G., Kirkegaard, J. B., Arosio, P., Michaels, T. C., Vendruscolo, M., Dobson, C. M., … & Knowles, T. P. (2016). Molecular mechanisms of protein aggregation from global fitting of kinetic models. Nature protocols, 11(2), 252–272.

[6] Mehta, P., Bukov, M., Wang, C.-H., Day, A. G. R., Richardson, C., Fisher, C. K., & Schwab, D. J. (2017). A high-bias, low-variance introduction to Machine Learning for physicists. Physics Reports, 810, 1–124.

[7] Raissi, M., Perdikaris, P., & Karniadakis, G. E. (2019). Physics-informed neural networks: A deep learning framework for solving forward and inverse problems involving nonlinear partial differential equations. Journal of Computational Physics, 378, 686–707.

[8] Meng, X., & Karniadakis, G. E. (2020). A composite neural network that learns from multi-fidelity data: Application to function approximation and inverse PDE problems. Journal of Computational Physics, 401, 109020.

[9] Karniadakis, G. E., Kevrekidis, I. G., Lu, L., Perdikaris, P., Wang, S., & Yang, L. (2021). Physics-informed machine learning. Nature Reviews Physics, 3(6), 422–440.

[10] Pateras, J., Rana, P., & Ghosh, P. (2023). A Taxonomic Survey of Physics-Informed Machine Learning. Applied Sciences, 13(12), 6892.

[11] Han, J., Jentzen, A., & E, W. (2018). Solving high-dimensional partial differential equations using deep learning. Proceedings of the National Academy of Sciences, 115(34), 8505–8510.

[12] Lu, L., Jin, P., Pang, G., Zhang, Z., & Karniadakis, G. E. (2021). Learning hidden physics of fluid flows. Science Advances, 7(15), eabc1515.

[13] Zhu, Y., Zabaras, N., Koutsourelakis, P. S., & Perdikaris, P. (2019). Physics-constrained deep learning for high-dimensional surrogate modeling and uncertainty quantification without labeled data. Journal of Computational Physics, 394, 56–81.

[14] Perdikaris, P., Raissi, M., Damianou, A., Lawrence, N. D., & Karniadakis, G. E. (2020). Nonlinear information fusion algorithms for data-efficient multi-fidelity modelling. Proceedings of the Royal Society A: Mathematical, Physical and Engineering Sciences, 476(2233), 20190436.

[15] Nasica-Labouze, J., Nguyen, P. H., Sterpone, F., Berthoumieu, O., Buchete, N. V., Cote, S., … & Derreumaux, P. (2015). Amyloid β protein and Alzheimer’s disease: When computer simulations complement experimental studies. Chemical reviews, 115(9), 3518–3563.

[16] Ghosh, P., Pateras, J., Rangachari, V., & Vaidya, A. (2021). A network thermodynamic analysis of amyloid aggregation along competing pathways. Applied mathematics and computation, 393, 125778.

[17] Dean, D. N., Das, P. K., Rana, P., Burg, F., Levites, Y., Morgan, S. E., … & Rangachari, V. (2017). Strain-specific fibril propagation by an Aβ dodecamer. Scientific reports, 7(1), 40787.

[18] Schaber, J. (2012). Easy parameter identifiability analysis with COPASI. Biosystems, 110(3), 183–185.

[19] Ghosh, P., Vaidya, A., Kumar, A., & Rangachari, V. (2016). Determination of critical nucleation number for a single nucleation amyloid-β aggregation model. Mathematical biosciences, 273, 70–79.

[20] Fawzi, N. L., Doucleff, M., Suh, J. Y., Clore, G. M. (2017). Mechanisms of Amyloid-Beta Protein Aggregation and Its Inhibition by Small Molecules. Biophysical Journal, 113(5), 1077–1088.

